# Auditory network persistence of stimulus representation in awake and naturally sleeping mice

**DOI:** 10.1101/2025.10.22.683945

**Authors:** Barak Hadad, Noa Regev, Uddi Kimchy, Ohad Rechnitz, Shai Abramson, Arseny Finkelstein, Dori Derdikman, Yuval Nir

## Abstract

Persistent neural activity often outlasts sensory stimulation, bridging perception and action. While commonly linked to working memory and decision making, its existence during passive states and sleep remains unclear. Using chronic high-density electrophysiology in freely behaving mice, we show that population spiking activity across the auditory cortical hierarchy enables decoding of past stimuli long after their offset, during both wakefulness and sleep. Time-resolved decoding revealed that in wakefulness, persistent representations decay uniformly across sensory and association cortices, whereas during sleep, persistence is prolonged in association cortex but remains brief in early auditory regions. Recurrent neural network modeling showed that higher internal noise during wakefulness reproduces this pattern, suggesting that reduced interference during sleep stabilizes sensory traces in associative areas. Our results demonstrate that persistent representation is a passive, state-dependent feature of sensory processing, supporting sensory maintenance even in the absence of active engagement.

## Introduction

When we fall asleep, we are partially disconnected from the environment and only rarely respond behaviorally to external stimuli. Yet, neurophysiological recordings have revealed that neurons in auditory cortex still respond robustly to sounds during both non-rapid eye movement (NREM) and REM sleep^1,2^. These responses exhibit altered dynamics compared to wakefulness, inducing post-stimulus silent (OFF) periods^3–5^ and reduced signatures of effective cortical connectivity^6^. The mechanisms underlying these changes remains unclear, with competing explanations including thalamic gating^7,8^ versus altered cortical network dynamics^9–11^.

One domain typically studied in the context of cognition during wakefulness is working memory - the ability to transiently maintain internal representations of recent sensory events^12–14^. Traditionally studied during delay periods in frontoparietal cortex under active (wake-attentive) task conditions, working memory is believed to bridge temporally coordinated behavior^15–17^. While sensory cortices have also been shown to exhibit stimulus-specific delayed activity^18–26^, these studies typically employ task-driven paradigms and behavioral engagement.

Here, we ask whether the neural representation of sensory information persists in the absence of task demands involving attention - both during passive wakefulness and during natural sleep. Using high-density electrophysiological recordings with Neuropixels probes in freely behaving mice, we simultaneously recorded spiking activity in hundreds of neurons across the auditory cortical hierarchy. Mainly, we recorded from primary auditory cortex (A1) and associative regions located ventrally or dorsally to A1 in temporal and parietal lobes. Polysomnographic monitoring enabled sleep scoring, allowing us to compare sensory persistence across well-defined vigilance states. By applying time-resolved population decoding, we assessed whether information about auditory stimuli could be reconstructed from neural activity after stimulus offset – potentially revealing a neural trace of the recent past.

Our results demonstrate that passive neural persistence of sensory information is a robust and widespread phenomenon: stimulus identity could be decoded across all states, often for as long as several seconds during sleep. These findings demonstrate that short-term sensory memory traces are maintained passively, even during sleep. We propose that these effects are population-level phenomena intrinsic to cortical processing irrespective of behavior, and are particularly apparent in sleep due to reduced influence of feedback connectivity, attention, or multisensory integration that may enhance the signal-to-noise ratios of persistent representation.

## Results

### Stable long-term recordings across auditory cortex during spontaneous sleep and wake

To investigate how recent sensory history is encoded across behavioral states, we recorded neuronal activity from freely moving mice across natural sleep-wake cycles. Adult male mice (C57BL/6JOlaHsd, age 10-12 weeks, N=5) were housed in their home cage placed inside an acoustic chamber and exposed to a stream of auditory stimuli while their brain activity was monitored (Methods, Figure 1A). Experiments started around 12PM, 2 hours after light onset, a time rich with NREM sleep and wakefulness periods. A custom 3D-printed head stage allowed stable recordings during unrestricted behavior (Figure 1B). Using high-density Neuropixels probes^27^, we simultaneously recorded from hundreds of neurons (n=178±65 per session, range: 93-277) spanning multiple levels of the auditory hierarchy, including primary auditory cortex (A1) and association regions located more ventrally (temporal association area, TeA, and perirhinal cortex, Prh) or dorsally (AUDd and somatosensory cortex) (Figure 1C).

**Figure 1.**
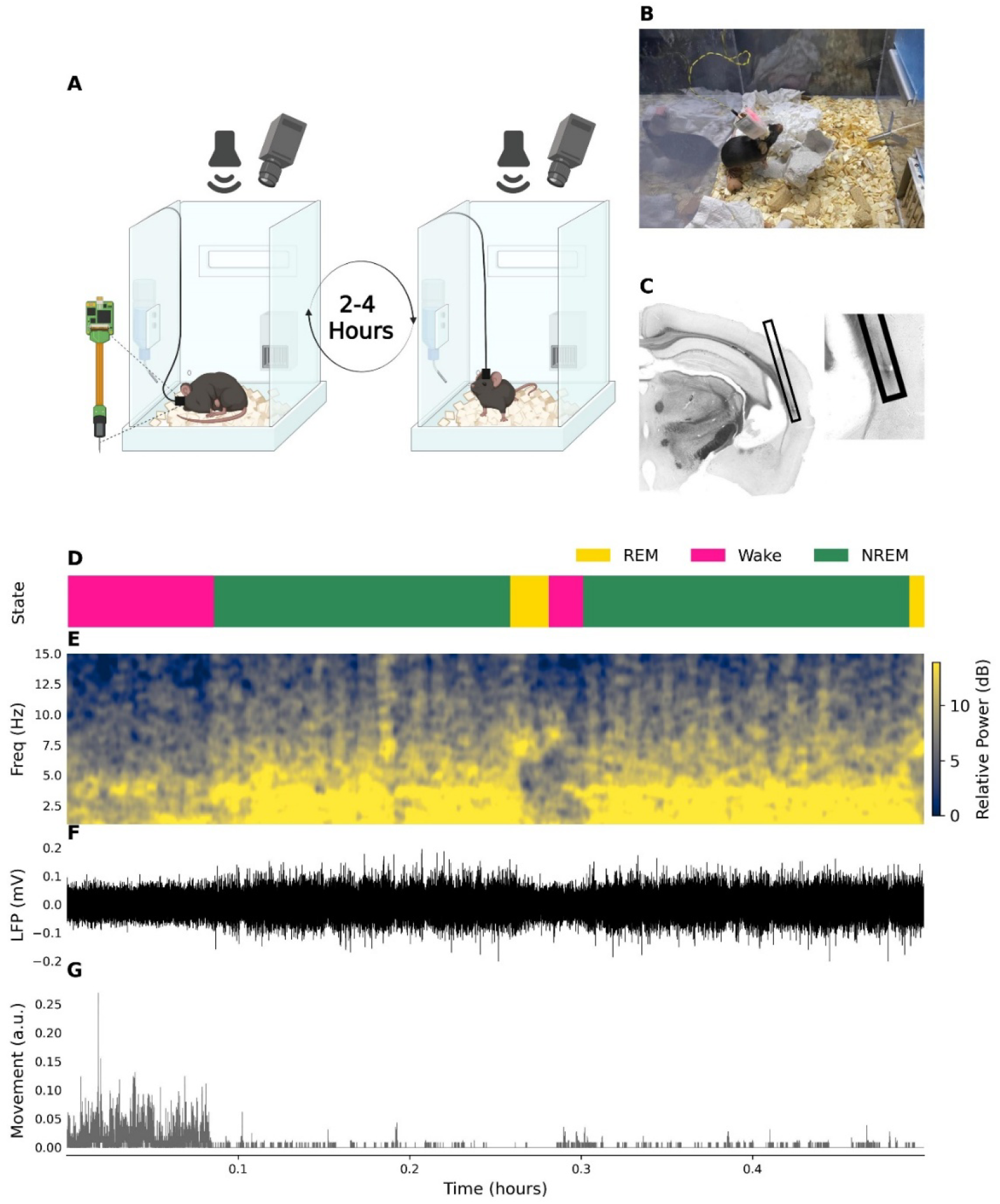
Experimental setup and behavioral state classification. (A) Schematic of the experimental setup. Mice were housed in their home cage within an acoustic chamber, equipped with an overhead ultrasonic speaker for stimulus delivery and a video camera for behavioral monitoring. Throughout the experiment, animals were allowed to sleep (left) or behave (right) freely while continuous extracellular neural activity was recorded (Created in https://BioRender.com). (B) Image of a freely moving mouse with a custom 3D-printed chronic head stage housing the Neuropixels probe. (C) Post-hoc histological validation of Neuropixels probe placement across the auditory cortical hierarchy. Probe trajectory marked with a black square. (D-G) Representative 30-minute segment of behavioral state classification using local field potential (LFP) features and movement tracking. Auditory stimuli were presented across all states. (D) State classification. Yellow, REM sleep; Pink, wake; Green, NREM sleep. (E) LFP spectrogram shows distinct spectral profiles for wake (mixed frequency activity), NREM (slow wave activity <4 [Hz]), and REM sleep (theta ~8 [Hz] activity). (F) Raw LFP trace. Note higher amplitude slow waves during NREM sleep. (G) Head movement metric from video monitoring used to detect behavioral activity.

Vigilance state classification (sleep/wake scoring) was achieved by combining local field potential (LFP) signals along with video-derived motion metrics^28,29^ (Figure 1D). Sleep stages were clearly distinguishable by characteristic changes in LFP spectral content and amplitude such as high-amplitude slow waves during NREM sleep or theta activity during REM sleep (Figure 1E-F), with movement traces further aiding the detection of wakefulness, its separation from REM sleep, and assisting in identifying transitions between states (Figure 1G). Due to the low proportion (<9%) of REM sleep in our data, we restricted our analysis to comparisons between NREM sleep and wakefulness.

Throughout the experiment, mice were passively exposed to three distinct auditory stimuli designed to engage different aspects of auditory processing: (1) a 40Hz click train, (2) dynamic random chords (DRC), and (3) broadband white noise (WN). All sounds were 2.5 seconds [s] in duration and presented in a random sequence, with silent intervals in between (average inter-stimulus-interval: 2.0 [s], with 0.5 [s] jitter). This design allowed us to compare spiking responses to identical sensory input across naturally occurring wakefulness and NREM sleep, and assess the generality of persistent representation across different sound statistics and cortical states.

### Auditory-evoked LFP responses vary across cortical hierarchy and behavioral state

To establish a systems-level view of auditory responses across the cortical hierarchy and identify early auditory vs. association cortices, we first examined LFPs evoked by a 40Hz click trains during wakefulness and NREM sleep states. Figure 2A shows the average evoked LFP response across all recording channels in one session, aligned to stimulus onset and offset (0 and 2.5 [s]). A strong, stimulus-locked response was evident throughout the sound presentation, most prominently in channels corresponding to primary auditory cortex (A1), as confirmed offline via histological reconstruction (Methods). Subsequent analyses focused on channels grouped into “Early” auditory regions (A1) vs. “Late” associative regions located more ventrally (TeA/Prh).

**Figure 2.**
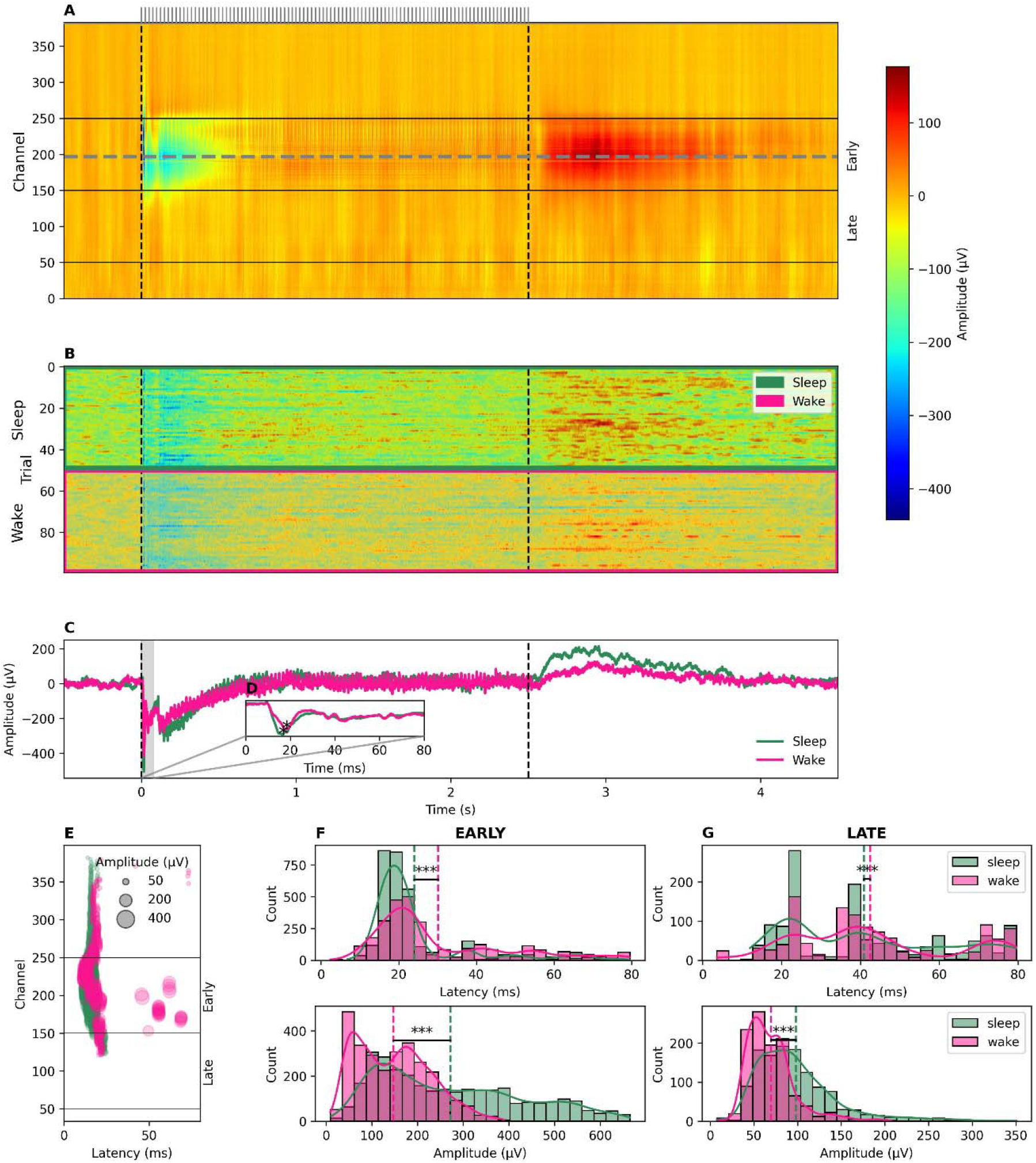
Representative Local field potential responses reveal state- and hierarchy-dependent auditory dynamics. (A) Average LFP response to 40 Hz click trains (averaged across n=100 trials) across all electrode channels spanning the auditory hierarchy. Dashed vertical lines indicate stimulus onset and offset (0 and 2.5 [s]). Robust phase-locked responses are observed during stimulus presentation, with the strongest responses localized to early regions (such as A1) as confirmed by histology. Channels are grouped into early and late regions. Dashed horizontal line indicates channel picked for panel B. (B) Trial-by-trial LFP traces from a representative channel (channel number 200) showing responses in 50 individual trials during wake (pink) and NREM sleep (green). As expected, sleep trials are associated with large-amplitude slow-wave activity whereas wake activity exhibits lower-amplitude faster dynamics. (C) Trial-averaged LFP responses in sleep and wake states. Sleep is associated with a larger onset (0-30 [ms]) negative response amplitude and a prominent positive offset rebound (~2.5-3.5 [s]). (D) Zoom-in on the initial 80 [ms] post-onset period, where latency and amplitude of the onset response minimum (asterisks) are extracted per state. (E) Latency and amplitude of the onset response across all channels. Each dot represents a channel; dot size reflects response amplitude. Early responding channels cluster in early auditory cortex with stronger and faster responses, while late regions show delayed, attenuated responses. (F-G) Early and late regions latency and amplitude of response across all animals and channels. Early regions were associated with significantly faster and stronger response compared to the late regions (p<0.001, via linear mixed model).

We next examined trial-level LFP responses from a representative channel with strong onset activity (Figure 2B). As expected, during NREM sleep, LFP traces showed larger amplitude slow waves and slower dynamics, while wake activity was faster and more temporally constrained. Trial-averaged responses (Figure 2C) further revealed that the LFP amplitude for the onset (0-80 [ms]) response was significantly higher (more negative) in sleep than in wake (p<0.001, via linear mixed model across all mice). Moreover, sleep was marked by a pronounced positive LFP deflection following stimulus offset, lasting ~1 second beyond sound termination - a pattern often associated with silent OFF periods^30^ and reduced in wakefulness.

To quantify the early sensory response, we extracted the latency and amplitude of the first post-onset minimum within the 0-80 [ms] window (Figure 2D, Methods). Extending this analysis across all channels, we found a hierarchical gradient in both metrics (Figure 2E). Across all mice, early auditory regions exhibited shorter LFP response latencies (sleep: 24.7±5.3 [ms], wake: 31.6±9.2 [ms]) and larger response amplitudes (sleep: −252.0±131.0 [uV], wake: −132.7±66.7 [uV]), while late areas showed slower (sleep: 45.2±12.3 [ms], wake: 48.6±8.9 [ms]) and weaker (sleep: −92.0±27.1 [uV], wake: −60.1±16.9 [uV]) responses (Figure 2F-G, p<0.001, via linear mixed model across all mice). These differences reflect both anatomical position and behavioral state in line with previous studies^3^, providing a basis for our subsequent analysis of persistent sensory traces across the cortical hierarchy.

### Spiking activity reveals state-dependent and hierarchical differences in auditory responsiveness

We next analyzed single-neuron spiking activity to characterize how auditory-evoked spiking responses vary across the auditory hierarchy and behavioral state. Neurons in early auditory regions (upper rows in Figure 3A) displayed clear, stimulus-locked responses across trials, whereas neurons in late regions (lower rows) exhibited more heterogeneous and temporally dispersed activity. Next, we inspected population activity by examining raster plots and peristimulus time histograms (PSTHs) in response to the three auditory stimuli. Representative examples show such responses for 40 Hz click trains (Figure 3B), dynamic random chords (Figure 3C), and white noise (Figure 3D). To further investigate this diversity at the single neuron level, we hand-picked four neurons that exhibited varying levels of responsiveness and stimuli selectivity (Figures 3E-G). These examples revealed that while each neuron maintained its unique response preferences across wake and sleep states, sleep consistently reduced firing rates during pre-, peri-, and post-stimulus intervals. A quantitative analysis across the entire dataset (n=1958 neuronal clusters) revealed that wakefulness was associated with higher spontaneous pre-stimulus firing rates (p<0.001, via linear mixed model), stronger responses during stimulus presentation (p<0.001, via linear mixed model), and higher spontaneous post-stimulus firing rates (p<0.001, via linear mixed model) than NREM sleep (Figure 3I).

**Figure 3.**
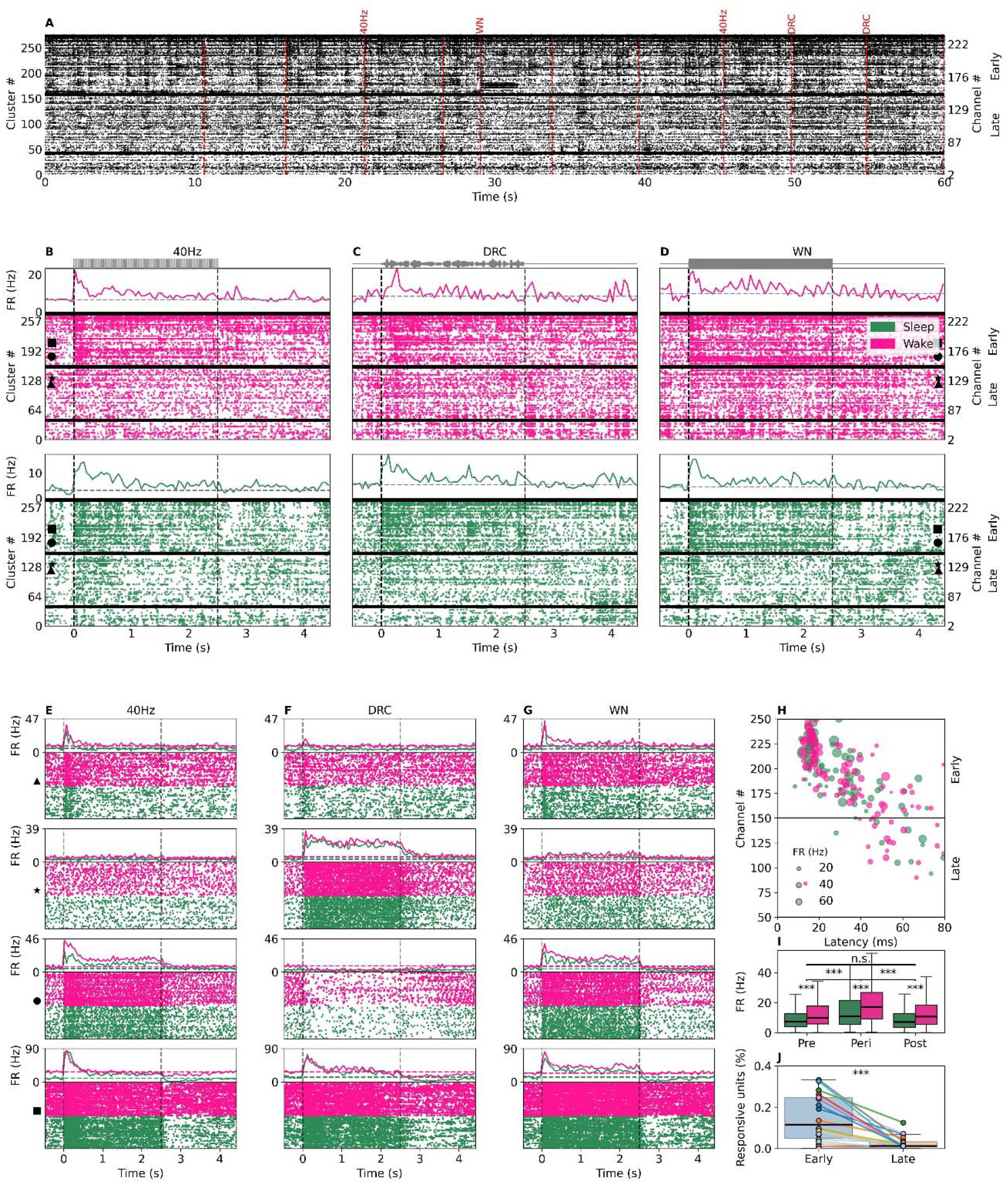
Spiking responses across the auditory hierarchy reveal state-dependent stimulus-specific dynamics. (A) Spiking activity of 277 recorded neurons over a 60-second period in one session, sorted by cluster identity (left) and recording depth (channel, right). Dashed vertical red lines mark auditory stimulus onsets. Note stimulus-locked responses in neurons from early auditory regions, while late auditory regions exhibit more variable patterns. (B-D) Raster plots (bottom) and population PSTHs (top) showing trial-averaged spiking activity during wake (pink) and NREM sleep (green) in response to 40 Hz click trains (B), dynamic random chords (C), and white noise (D). Neurons are sorted as in panel A. (E-G) Example single-unit responses illustrating the diversity in tuning and temporal dynamics (e.g. neuron in second row prefers DRC stimulus while neuron in third row prefers anything but DRCs). Raster plots show 50 trials per sound type (wake: pink; sleep: green); top traces show PSTHs across all stimuli. (E) 4 single-unit responses to 40 Hz click trains. (F) 4 single-unit responses to DRC stimuli. (G) 4 single-unit responses to white noise stimuli. Note similar responses across wakefulness and NREM sleep despite overall higher firing rates in wake across pre-, peri-, and post-stimulus periods. (H) Characterization of onset response latency and amplitude across all responsive neurons (Methods). Dot size reflects response amplitude. Neurons in early auditory regions show shorter latencies (typically < 20 [ms]) and higher amplitudes than late auditory areas. (I) Firing rate analysis across behavioral states revealed consistently elevated neural activity during wakefulness compared to NREM sleep (N=5 mice). This state-dependent modulation was observed across all temporal windows-pre-, peri-, and post-stimulus periods. No significant differences between pre- and post-stimulus firing rates within each state. (J) Regional analysis of stimulus responsiveness demonstrated a hierarchical gradient in auditory processing. Early auditory regions exhibited significantly higher proportions of responsive units compared to late auditory areas (15.2±11.6% vs. 2.5±3.1%, p<0.001, Wilcoxon signed-rank test), with each data point representing the percentage of responsive neurons per recording session.

Next, to compare between early and late auditory regions, we quantified the latency and amplitude of each neuron’s initial onset response (Figure 3H). Consistent with the LFP results, neurons in early auditory regions responded faster (p<0.001, via linear mixed model) and with higher amplitude (p<0.01, via linear mixed model) than those in late regions. Furthermore, we observed a higher density of stimulus-onset responsive neurons in early regions (15.2±11.6% in early vs. 2.5±3.1% in late, p<0.001, via Wilcoxon signed-rank test) suggesting a hierarchical gradient in auditory encoding precision associated with more distributed representation in early regions vs sparser representations in late regions (Figure 3J).

### Persistent post-stimulus decoding of stimulus identity from population activity

To test whether sensory representations persist beyond stimulus offset, we trained a logistic regression-based classifier to decode stimulus identity from population spiking activity across cortical regions and behavioral states (Methods, Figure 4A). Separate models were trained for binary classification (a specific sound vs. other options) and evaluated using either spiking activities in early or late auditory regions, and separately using either wake or NREM sleep data. This allowed us to assess not only stimulus encoding, but also whether decoding accuracy persisted after sound termination.

**Figure 4.**
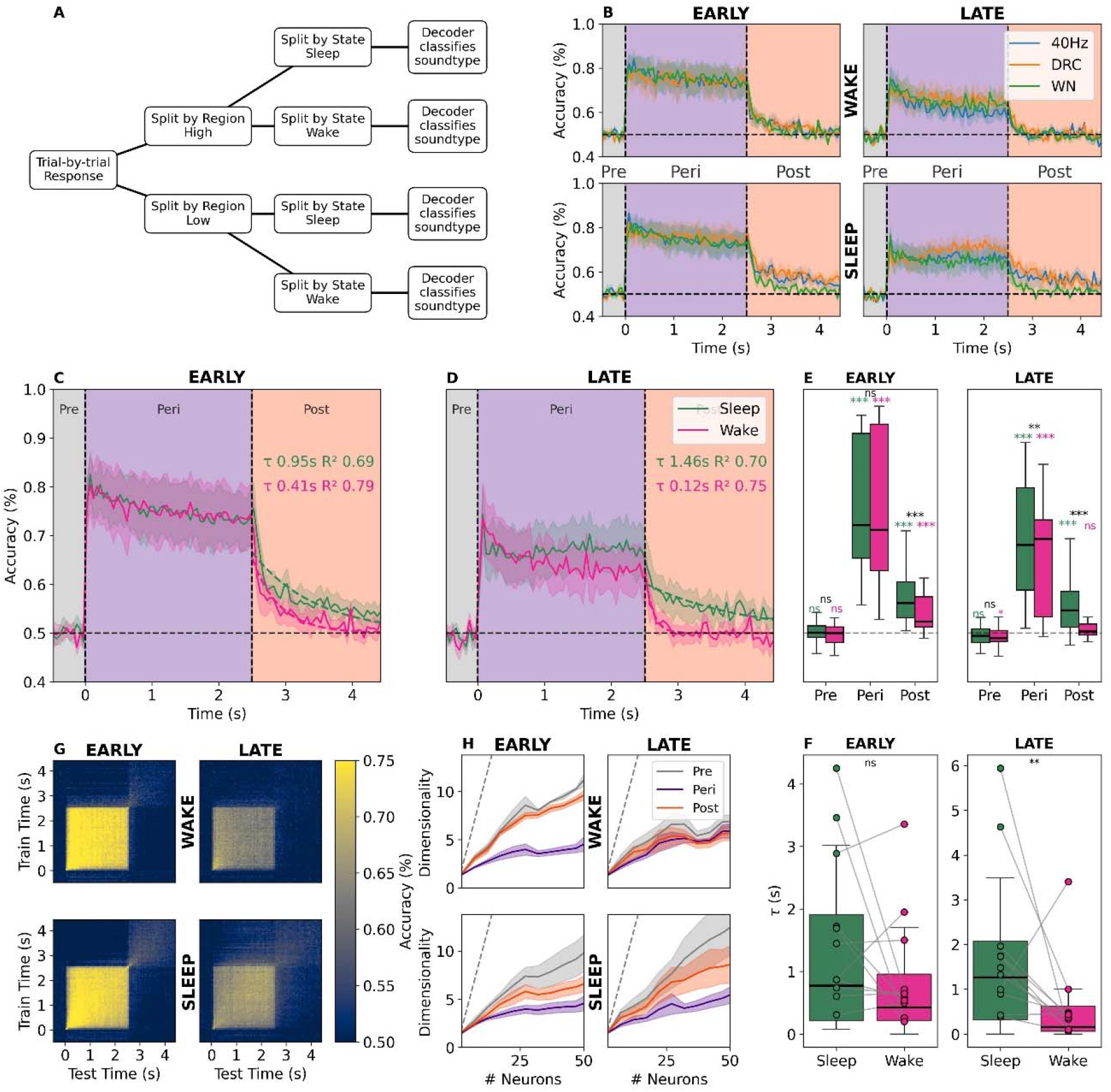
Logistic regression decoding reveals stimulus-specific representation persisting after sound offset. (A) Schematic of the decoding pipeline. Neuronal population responses were grouped by cortical region and by behavioral state. For each condition (region, state), a logistic regression decoder was trained in 50 [ms] bins to classify stimulus identity in a binary manner (that specific stimulus vs. other options) based on trial-by-trial population spiking. (B) Dynamics of classification accuracy for each region (columns) and state (rows), across the three stimulus types (40 Hz click, DRC, white noise). During stimulus presentation, decoding accuracy was robust in both regions and states. While accuracy was at chance (50%) before stimulus onset (pre-stimulus baseline), it remained significantly above chance post-stimulus offset - particularly in early areas and during sleep, with significant decoding persisting for seconds. (C-D) Mean decoding accuracy across stimuli, plotted as a function of time for early (C) and late (D) auditory regions. Exponential fits to the post-stimulus decay revealed similar decay constants 0.41 [s] for wake and 0.95 [s] for sleep in the early region, but markedly longer persistence in late regions during sleep (1.46 [s] vs. 0.12 [s] in wake). (E) Regional comparison of decoding performance revealed distinct temporal profiles across the auditory hierarchy. In early auditory areas, classification accuracy remained significantly above chance during both peri- and post-stimulus periods, with sleep exhibiting significantly higher accuracy than wakefulness specifically during the post-stimulus window. In late auditory regions, both sleep and wake conditions showed above-chance accuracy during stimulus presentation, but only sleep maintained significant decoding performance in the post-stimulus period. All tests were performed via linear mixed model. (F) Quantitative analysis of decay dynamics confirmed regional and state-dependent differences in representation persistence. Early auditory regions exhibited similar decay constants between wake and sleep conditions (no significant difference), while late auditory regions demonstrated enhanced stimulus representation persistence during NREM sleep. (G) Temporal generalization analysis. Performance accuracy for classifiers trained on each time point (y-axis) vs. tested on every other time point (x-axis). High decoding generalization was confined to the stimulus duration window (clear squares), but traces of post-stimulus generalization emerge in early regions and in sleep. (H) Dimensionality of the neural population response during pre-(gray), peri-(purple), and post-stimulus (orange) windows. Dimensionality increases least during the peri-stimulus window, and remains lower than pre-during the post-stimulus window - particularly in sleep and early regions. In late regions, during wakefulness, hardly any increase was observed.

During stimulus presentation, all models achieved robust above-chance classification accuracy, confirming that auditory classification information is encoded in the neural population (Figure 4B). Decoding accuracy was highest in the early auditory cortex (~80%) and slightly reduced in late areas (~70%) across both states (Figures 4C-D), likely reflecting multiple driving factors including differences in selectivity and number of responsive neurons. As expected, decoding accuracy was at chance prior to sound onset and significantly above chance during sound presentation for both regions and states. Crucially, however, significant decoding remained above chance for several hundred milliseconds after stimulus offset. For the early region, for both sleep and wake versus chance, results were significant (p<0.001, via linear mixed model). On the other hand, for the late region, wake against chance was not significant while for sleep it was (p<0.001 via linear mixed model). In both regions, sleep against wake showed a significant difference (p<0.001 via a linear mixed model) with sleep accuracy being higher than wake (Figure 4E).

To quantify the temporal dynamics of this persistent representation, we computed the average decoding accuracy across all stimulus types and fit exponential decay curves to the post-stimulus period. The persistence varied by both region and state: early regions showed similar decay dynamics in wake and sleep, while late areas, during NREM sleep, exhibited prolonged persistence. In early regions, decay time constants were equivalent in wake and sleep (no significant difference, see Figure 4F, “early” panel). In contrast, late areas showed a pronounced state difference: persistence decayed more than ten times slower in sleep than in wake (Figure 4F, “late” panel). This finding suggests a sleep-specific enhancement of post-stimulus retention in associative areas.

To further explore the structure of this persistence, we performed temporal generalization analysis - training the decoder on a single time point and testing on all others (Figure 4G). A strong square-shaped off-diagonal emerged during stimulus presentation in all conditions, indicating temporally consistent encoding. After stimulus offset, generalization faded, but significant residual accuracy remained evident - particularly in the early region and during sleep. Notably, late areas during wake exhibited only modest post-stimulus generalization, suggesting that representation persistence is both anatomically- and state-specific.

Finally, we examined the dimensionality of the neural response during different time windows (Figure 4H). As expected, the stimulus window exhibited the lowest dimensionality, likely reflecting the fact that all neurons are locked to a single driving stimuli^31,32^. The post-stimulus window showed intermediate dimensionality - lower than pre-stimulus activity, but higher than during sound - consistent with a structured but decaying persistent activity. This pattern held across most conditions, with the notable exception of late areas during wake, where no post-stimulus dimensionality increase was observed.

Together, these results reveal that auditory representations persist beyond stimulus offset in both wake and sleep. Persistent representation is strongest in early areas and is selectively enhanced during sleep in late regions - suggesting a state-dependent shift in the temporal integration of sensory information.

### A minimal RNN model reproduces state- and region-specific persistence through noise modulation

To probe how circuit level mechanisms could account for the persistent representations observed experimentally and their differential expression across early versus late auditory regions and across states, we first classified neurons into two groups: those showing a significant difference in firing rate during the post stimulus period relative to pre stimulus, and all others (Figure 5A). We refer to the former as “persistent units,” consistent with prior literature on sustained activity in working memory tasks^15,19,23^. The fraction of persistent units varied across both state and region (Figure 5B–C). In both sleep and wake, persistent units were more prevalent in early regions (sleep: 44.7±20.8%; wake: 34.8±11.9%) than in late regions (sleep: 30.6±12.0%; wake: 25.0±5.9%). To test whether decoding persistence was driven by these units, we removed them from the decoding pipeline. Remarkably, the decoding accuracy trajectory still exhibited robust persistence (Figure 5D–E), demonstrating that population level persistence is not reducible to the activity of a small subset of persistent neurons.

**Figure 5.**
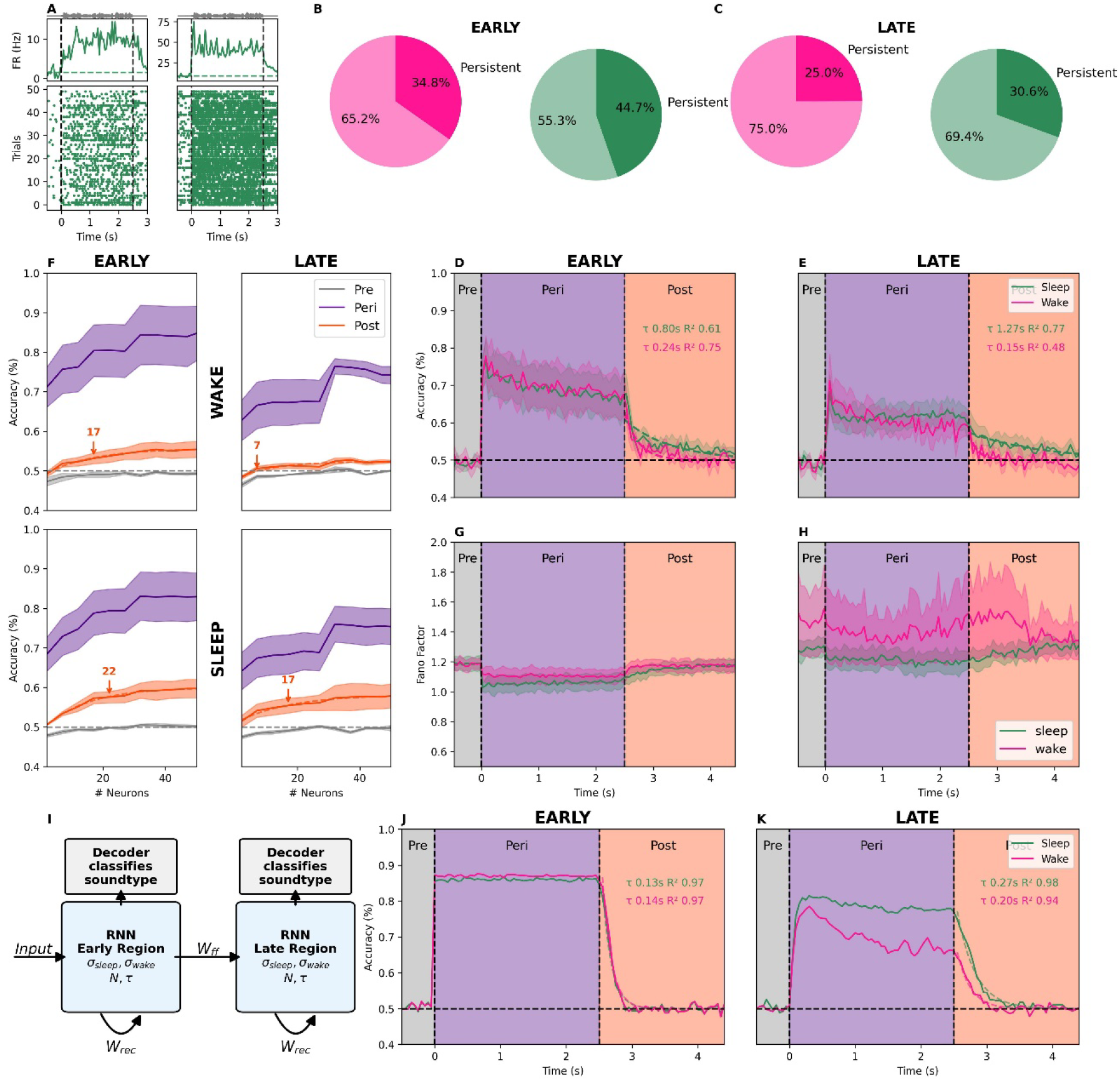
A recurrent neural network (RNN) model reproduces hierarchical and state-dependent persistence through noise and recurrence. (A) Two example persistent units showing significant changes in firing rate at post-stimulus time window relative to the pre stimulus baseline. Top: PSTH showing the mean firing rate across time, Bottom: Raster plot showing all spikes. (B–C) Fraction of persistent clusters across states and regions. (D–E) Decoding accuracy after removing persistent clusters, showing that persistent representation remains even without these units. (F) Decoding accuracy as a function of neuron number during pre-(gray), peri-(purple), and post-stimulus (orange) windows. Accuracy improves with population size and plateaus at a condition specific threshold (arrows). (G–H) Fano factors for early and late regions across states. (I) Schematic of a two layer RNN representing early and late auditory regions, each with 22 recurrent units. The integration constant (*τ*) is fixed, while internal noise (σ) is tuned separately for state and region based on the Fano factors. (J) Decoding from the simulated early population replicates experimental persistence with similar decay in sleep and wake. (K) Decoding from the simulated late population shows reduced accuracy during stimulation and state dependent divergence post stimulus, with rapid decay in wake and sustained persistence in sleep.

We next asked whether persistence reflects a distributed population phenomenon. By systematically increasing the number of neurons included in decoding, ranked by decoder weights, we found that post-stimulus decoding accuracy increased with population size and plateaued after approximately 7-22 clusters depending on state and region (Figure 5F; see Methods). This result indicates that persistent sensory representation emerges from collective activity across a distributed ensemble rather than from a few strongly active units. Importantly, this effect was absent in pre stimulus windows, underscoring that persistence depends on coordinated post stimulus activity across the population.

To examine whether differences in variability and noise might underlie state and region dependent dynamics, we quantified Fano factors per cluster across states and regions (Figure 5G–H). In early regions, Fano factors were comparable between sleep and wake. In late regions, Fano factors were elevated relative to early regions, with a striking divergence across states. While sleep values followed a similar trend to early regions, albeit higher overall, wake showed substantially greater variability. Across all regions and states, variability was quenched after stimulus onset^32^, except in late regions during wake, where variability remained high. This exception may explain the absence of persistent representation in that condition.

Motivated by these findings, we developed a minimal two-layer recurrent neural network (RNN) model to capture how changes in internal noise could give rise to the observed state and region specific persistence (Figure 5I). The first layer, receiving external input, represented early auditory cortex, while the second layer, driven by feedforward input, represented late regions.

Each layer comprised 22 recurrently connected units, consistent with our empirical observation that decoding accuracy saturates around this population size (Figure 5F).

We implemented each layer as a discrete time dynamical system (see Methods), where activity at time t depended on external input, recurrent dynamics, and feedforward input. Two key parameters governed each layer: the recurrent integration constant (*τ*, leak term), and Gaussian noise amplitude (σ). Crucially, the noise parameter was not set arbitrarily but was tuned directly from the Fano factors measured in the data. We extracted Fano factor values separately for pre-, peri-, and post-stimulus windows in each state and region, and separately for individual neurons (Figure 5G-H) and for the neuronal population and used these empirical values to modulate the internal noise level of the model dynamically (Methods).

Simulations of spiking population responses to noisy stimuli reproduced the main experimental findings. In the early layer, decoding rose during stimulus presentation and persisted after offset, with comparable decay rates across sleep and wake (Figure 5J). In the late layer, decoding accuracy during the stimulus was lower, and post stimulus persistence diverged across states. Persistence was sustained in sleep for several seconds, but decayed rapidly in wake (Figure 5K). These patterns closely matched empirical recordings.

The model therefore links state dependent persistence to differences in intrinsic noise. Elevated variability during wake, likely reflecting ongoing attention, multisensory integration, and motor related activity, degrades persistence. Reduced noise during sleep enables robust post-stimulus decodability. Importantly, this match to experimental data emerged solely from incorporating empirically measured noise levels across time windows. Together, these results demonstrate that state and region specific persistence can emerge from basic circuit level properties without invoking top down memory mechanisms. A simple reduction in internal noise during sleep is sufficient to sustain stimulus representations beyond offset^33,34^.

## Discussion

To what extent does cortical spiking activity represent information about the immediate past when an event is no longer present in the environment? Traditionally, such questions have been explored through the lens of working memory and active task engagement, with persistent neural activity viewed as reflecting a cognitive process primarily occurring in prefrontal or parietal circuits during delay periods. In this study, we demonstrate that some persistent representation of sensory input emerges passively in the auditory cortex, even in the absence of task demands or conscious awareness (NREM sleep). Our findings reveal that neural representations of sounds persist beyond stimulus offset during both wakefulness and NREM sleep, with distinct dynamics across the auditory cortical hierarchy and behavioral states. Thus, persistent activity - a core feature of working memory – may be driven in part by passive intrinsic features of information processing irrespective of cognition.

### Persistence of sensory representation across states

Using large-scale electrophysiology and model-based decoding, we show that stimulus-specific representation remains decodable from population spiking activity, well beyond stimulus termination. This persistent representation does not necessarily require top-down engagement or behavioral relevance, but emerges passively as a property of sensory processing. These memory-like traces are present not only in the awake brain but also during NREM sleep - a state in which the brain is considered to be relatively disconnected from the environment. Importantly, this post-stimulus persistence of representation was observed across different stimulus types (Figure 4B), including white noise, click trains, and dynamic random chords, highlighting the generality of this phenomenon.

These results blur the classical distinction between attention-dependent working memory and task-free sensory processing. Persistent activity has previously been reported in primary visual^18,22^, somatosensory^19,35^, and auditory cortices^23,24,26,36^ during memory tasks, and even in spiking activity of individual human neurons^37^. Our findings extend these observations to conditions without explicit behavioral demands, including during sleep. This suggests that persistence can arise from network dynamics, rather than requiring top-down attention.

### State- and region-specific dynamics of persistent representation

By analyzing the time course of decoding accuracy across brain regions and states, we demonstrated an interaction between behavioral state and cortical hierarchy. In early auditory regions, the temporal profile of persistence was comparable between wakefulness and NREM sleep, with no significant difference in post-stimulus decay constants. In contrast, late regions exhibited strong state-dependent dynamics. During NREM sleep, decoding accuracy decayed significantly more slowly than during wake.

At this point we can only speculate about the source of longer persistent representation in late areas during NREM sleep compared to that observed in early regions. One possibility is that during wakefulness, multiple processes such as effects of motor actions^34^, multi-sensory integration^10^, prediction and top-down feedback^6,38^, interfere with the persistent representation. By contrast, such processes may be less prominent during NREM sleep, allowing associative areas to maintain stimulus-related information for longer periods of time.

### Neural population-level correlates of post-stimulus persistence

Importantly, our results also clarify the population-level structure of representation persistence. We find that post-stimulus decoding accuracy depends on the number of neurons included in the model, saturating after 22 neurons - consistent with a distributed coding scheme. Furthermore, dimensionality analyses revealed that during stimulus presentation, the response space becomes more low dimensional than during pre-stimulus - likely reflecting stimulus-driven coordination^32^. The dimensionality of the post-stimulus window is intermediate, suggesting that persistence arises from a structured but decaying representation of past input.

These results point toward a view of persistence not as a static echo, but as a dynamic, structured transition in the neural state space. That this structure survives well into NREM sleep implies that even in a state classically defined by sensory disconnection, the brain maintains an internal model of recent events.

### Mechanistic insights from a minimal recurrent network model

To determine whether the observed dynamics could arise from simple network-level mechanisms, we developed a two-layer recurrent neural network model reflecting the hierarchical organization of auditory cortex. By only adjusting the internal noise (higher during wake) - we were able to reproduce the key features of our data: similar persistence in early regions across states, and enhanced persistence in late areas during sleep.

This result supports the hypothesis that some persistent neural activity may not require specialized memory circuits or top-down modulation. Instead, the temporal retention of sensory information may emerge naturally from basic principles of circuit organization: recurrent connectivity, noise levels, and feedforward coupling. These findings resonate with recent theories proposing that memory traces are distributed across sensory hierarchies and can emerge from spontaneous population dynamics^39,40^.

### Limitations

While our findings reveal robust, state- and region-dependent representation persistence, several limitations warrant consideration. First, our recordings were limited to auditory presentation in freely behaving mice without an explicit task. It remains unknown whether similar persistence dynamics occur in other sensory modalities during passive and offline states. Second, our analysis focused on NREM sleep, as REM sleep episodes were infrequent and variable. Given that REM sleep is characterized by different oscillatory and neurochemical dynamics^41^, future studies are needed to determine whether the patterns we report generalize to REM sleep. Third, our analyses and minimal RNN model capture persistence through recurrence and noise modulation but do not address more complex circuit motifs or cell-type specific contributions. Future studies are needed to shed additional light on the cellular and circuit mechanisms driving passive persistent activity. Finally, although beyond the scope of this work, we cannot rule out acoustic confounds contributing to response variability in wakefulness.

Mice can reorient their ears during wake but not sleep, potentially altering sound features and affecting persistent representations as the features of auditory stimuli change.

### Implications for sleep, memory, and cognition

Together, our findings suggest that persistence of sensory representations is a fundamental property of cortical processing, rather than an exclusive signature of conscious task-related cognition. The fact that memory-like traces persist in sleep, and do so more robustly in late areas, raises new questions about the role of sleep in passive sensory integration and early memory formation. A key question for future studies is whether the existence of persistent sensory activity we observe here during sleep is associated with subsequent memory and behavioral changes, as is the case for example with targeted memory reactivation^42^ (TMR). If so, above and beyond consolidating memories encoded in wakefulness, the sleeping brain may also retain and structure passively encountered sensory events affecting future behaviors. How these persistent traces are relayed and integrated in hippocampal–entorhinal circuits remain unclear, and further research could investigate the pathways through which cortical activity contributes to offline memory processing.

Altogether, these findings open new avenues for research into sleep-dependent plasticity, attentional modulation, and the balance between task-free and task-driven memory processes. They also point toward the use of decoding-based, model-driven population analyses as a powerful framework for understanding how the brain maintains information over time and across states.

## Methods

### Animals

All experimental procedures including animal handling, surgery, and experiments followed the NIH Guide for the Care and Use of Laboratory Animals and were approved by the Institutional Animal Care and Use Committee (IACUC) of Tel Aviv University (approval 2409 - 165 - 5). Electrophysiological recordings were conducted in five adult male mice (strain: [C57BL/6JOlaHsd], age: [10-12 weeks], weight: [29-33gr]). Mice were housed individually in transparent Perspex cages under controlled ambient conditions (temperature: 20-24 °C) with ad libitum access to food and water. The housing environment was maintained on a 12 h light/dark cycle with lights on at 10:00 AM.

### Surgical Procedures and Electrophysiological Recordings

Mice were chronically implanted with Neuropixels 1.0 probes^27^ targeting the auditory cortical hierarchy. Prior to surgery, the probes were coated with the fluorescent lipophilic dye DiIC18 (D3911; Rhenium) to aid post hoc histological localization. General anesthesia was induced with isoflurane (4%) and maintained at 1.5-2% throughout the surgery. Mice were secured in a stereotaxic frame (David Kopf Instruments), and body temperature was maintained at 37 °C using a closed-loop heating system (Harvard Apparatus).

Animals received preoperative treatments including Buprenorphine (0.05 mg/kg, s.c.) and carprofen (5 mg/kg, i.p.). Eyes were protected with Viscotears gel, and local anesthesia was administered via subcutaneous lidocaine (3 mg/kg) before incision. The scalp was shaved and disinfected, and a small craniotomy was made over the right auditory cortex. The dura was carefully removed, and probes were stereotactically inserted at a 12° angle to traverse multiple regions along the auditory hierarchy, from primary auditory cortex (A1) through secondary areas (e.g., TeA) to higher-order associative regions (e.g., perirhinal cortex). Coordinates relative to bregma were anteroposterior (AP): −2.5 mm; medio-lateral (ML): 4.0 mm, dorso-ventral (DV): − 3.2 mm relative to the brain surface.

A custom 3D-printed enclosure protected the probe and allowed free mouse movement within the home cage. The craniotomy was sealed with silicone gel (Kwik-Sil, World Precision Instruments), and the probe was secured with dental cement (Fusio, Pentron). Post-surgery, mice received systemic buprenorphine (0.05 mg/kg, s.c.) for analgesia. A bright green marker was affixed to the head stage to facilitate subsequent video analysis.

Neural signals were continuously recorded at 30⍰kHz. Local field potentials (LFPs) and spike data were processed separately. Spike sorting was performed using Kilosort 4.0^43^, and units were automatically and manually curated based on waveform characteristics and refractory period violations using Bombcell^44^.

### Histological Verification

Following the completion of recordings, electrode placement was verified histologically (see Figure11C). Mice were deeply anesthetized with isoflurane (5%) and transcardially perfused with 4% paraformaldehyde (PFA) in phosphate-buffered saline (PBS). Brains were post-fixed in 4% PFA at 4° C for one week before being sectioned into 50-60 μm coronal slices using a vibrating microtome (Leica Biosystems). Sections were stained with fluorescent cresyl violet/Nissl stain (Rhenium) and imaged using fluorescence microscopy to identify DiI-labeled probe tracks. Recording site positions were matched to anatomical landmarks and registered to the Allen Mouse Brain Atlas to assign neurons to specific cortical and subcortical regions along the auditory hierarchy.

### Experimental Design

Following surgical recovery, mice were individually housed in their home cages (see Figure11A and 1B) set inside a double-wall sound-attenuating acoustic chamber (Industrial Acoustics Company, Winchester, UK) to minimize environmental noise and reverberations^45^. During this time, animals were gradually acclimated to the experimental setup, including habituation to auditory stimulation, through gradually increasing durations of passive auditory exposure (ramping up from 30 minutes to 3 hours over days). Experimental sessions started after at least a week post-surgery. Each mouse participated in 2-3 recording sessions, during which auditory stimuli were continuously presented for 2-4 hours while the animal was freely behaving in its home cage. Stimuli were delivered throughout the sessions, which included spontaneously occurring wakefulness and sleep, independent of behavioral state.

### Auditory Stimulation

Auditory stimuli were generated using MATLAB (MathWorks) and converted to analog voltage signals via a high-fidelity sound card (192 kHz; LynxTWO, Lynx Studio Technology). Signals were amplified (SA1, Tucker-Davis Technologies) and delivered through a magnetic speaker (MF1, TDT) mounted 60 cm above the animal’s home cage. The sound pressure level at the cage floor was calibrated to ~70 dB SPL using a measurement microphone (DVM805, Velleman). Stimuli included: 40 Hz click trains, Single click pulses, Dynamic Random Chords (DRCs), Broadband white noise bursts, three distinct ultrasonic vocalizations (USVs). For consistency in time-resolved and decoding analyses, we focused on three stimuli with identical 2.5-second-long duration - the Dynamic Random Chords (DRC), 40 Hz click trains, and white noise bursts - which permitted direct comparison of stimulus-driven and post-stimulus activity dynamics. Stimulus delivery occurred during 2-4 hour long sessions of continuous recording. Each hour contained approximately 100 repetitions per stimulus type, presented in random order. The inter-stimulus interval was 2.0 ± 0.5 s, with an extended 6-second pause inserted every 10 minutes to allow for recovery from potential adaptation effects. Stimuli were presented passively throughout both wake and sleep epochs, following two days of habituation during which mice were exposed to the full stimulus set in the same environment without concurrent recordings.

### Sleep Scoring and Polysomnography

Vigilance states were classified using a combination of electrophysiological and behavioral features, integrating local field potential (LFP) spectral and temporal dynamics with a head movement metric. Head movement was quantified via a computer vision algorithm that tracked the center of mass of a bright green marker affixed to the head stage. Frame-by-frame positional changes were computed to yield a continuous motion trace. LFP signals were down-sampled, bandpass filtered, and analyzed using short-time Fourier transforms to generate continuous spectrograms. Wakefulness, NREM sleep, and REM sleep were manually scored by an expert blinded to auditory stimulation protocol. Scoring was based on standard polysomnographic criteria, including delta/theta power ratios, the presence of slow-wave activity, and behavioral quiescence (i.e., sustained absence of movement). Because REM sleep occurred infrequently in our dataset, all subsequent analyses focused exclusively on wake and NREM trials. This ensured adequate statistical power and consistent comparisons across states.

### Preprocessing and Neuronal Inclusion Criteria

Spike sorting was performed using Kilosort 4.0^43^ and automatically curated using Bombcell^44^, followed by manual verification and curation with Phy^46^, to separate high-quality single units and multi-unit activity (MUA) from noise clusters. Clusters were anatomically categorized into early and late auditory regions based on their relative depth along the Neuropixels probe and confirmed by histological reconstruction. Across five animals, the average number of well-isolated clusters per session was 61±34 in the early regions and 65±35 in the late regions (mean ± standard deviation).

### Latency and amplitude characterization

Latencies and amplitudes were computed using a cluster-based permutation test to identify periods of significant deviation from baseline activity. For each unit or channel, we first compared the time-resolved signal to its pre-stimulus baseline across trials, identifying contiguous clusters of time points showing statistically significant changes. The latency was defined as the onset of the earliest significant cluster, capturing the initial response to the stimulus. Within this cluster, the peak value was used to determine both the response amplitude and the precise latency of maximal activation.

### Decoding Analysis

To assess stimulus-specific neural representations, we trained a logistic regression decoder to decode stimulus identity based on population spiking activity. Spikes were binned in 50-ms non-overlapping windows and used as input features. Separate decoders were trained for each cortical region (early-order vs. late-order), brain state (wake vs. NREM), and stimulus type (401Hz click train, DRC, white noise). Each model was trained to classify whether a given stimulus was being played (vs. others) during a specific time bin such that chance performance is 50%. Models were trained on 80% of trials and tested on the remaining 20% (cross-validated). For each configuration, models were trained on individual time bins and tested across all time bins to assess temporal generalizability of the population code. Decoding performance was quantified as classification accuracy. To quantify persistence after stimulus offset, we computed decoding accuracy in post-stimulus bins and fitted exponential decay functions of the form: 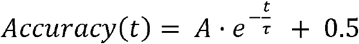 to estimate temporal decay constants (*τ*). The final decoding curves per condition represent the average of individual sound-specific models (one-vs-all classification), providing a robust estimate of generalizable stimulus encoding and persistence.

### Population Size Analysis

To evaluate how decoding accuracy scales with neuronal ensemble size, we conducted a cumulative inclusion analysis based on decoder feature importance. Neurons were rank-ordered by the absolute magnitude of their decoder weights during the stimulus window, reflecting their individual contributions to stimulus classification. We incrementally added neurons to the decoding model in order of decreasing importance and recorded the resulting classification accuracy. This procedure was performed independently for each brain state (wake vs. NREM), cortical region (early vs. late), and time window of interest: pre-, peri-, and post-stimulus. As expected, decoding accuracy during stimulus presentation improved as more informative neurons were added. Post-stimulus decoding, reflecting persistent activity, also improved with population size, though the effect varied by region and state. In contrast, decoding during pre-stimulus spontaneous activity remained at chance, demonstrating the specificity of stimulus-related coding.

To quantify the plateau in post-stimulus decoding, we fitted a saturating exponential function to the accuracy as a function of neuron number, 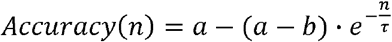, where *a* represents the asymptotic maximum decoding accuracy, *b* is the starting accuracy with zero neurons, *n* is the number of neurons included, and *τ* is the rate constant governing how quickly accuracy approaches the asymptote. The plateau ensemble size was defined as the number of neurons required to reach 95% of the asymptotic value *a*.

### Dimensionality Analysis

To assess the complexity of population-level neural responses, we computed the effective dimensionality of binned firing rate activity using principal component analysis (PCA). Analyses were performed separately for each time epoch: spontaneous (pre-stimulus baseline), stimulus (during sound presentation), and persistent (post-stimulus period), across both brain states (wake vs. NREM) and cortical regions (early vs. late). Dimensionality was quantified using the participation ratio of the PCA eigenvalue spectrum, a well-established metric for estimating the effective number of independent dimensions contributing to population activity^31,32,47^:

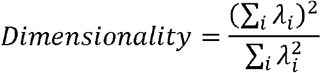

Where *λ*_*I*_ denotes the variance explained by the *i*th principal component.

This measure captures how distributed or concentrated the population response is across principal axes. To examine how dimensionality scales with population size, neurons were not randomly sampled; instead, they were rank-ordered based on their decoder coefficients during the stimulus window, prioritizing those that contributed most to decoding accuracy. Neurons were then added sequentially according to this ranking, from most to least informative. For each incremental population size (ranging from 2 to 50 units), we computed the dimensionality and compared it to the identity line, which reflects the dimensionality of a fully uncorrelated system.

### Recurrent Neural Network (RNN) Simulations

To model the temporal evolution of sensory responses across cortical hierarchies and brain states, we implemented a minimal rate-based recurrent neural network (RNN) with two layers representing early (E) and late (L) auditory regions. Each layer comprised 22 neurons with discrete-time dynamics governed by recurrent inputs, feedforward coupling, stimulus drive, and Gaussian noise.

The network dynamics were defined by the following update equations:

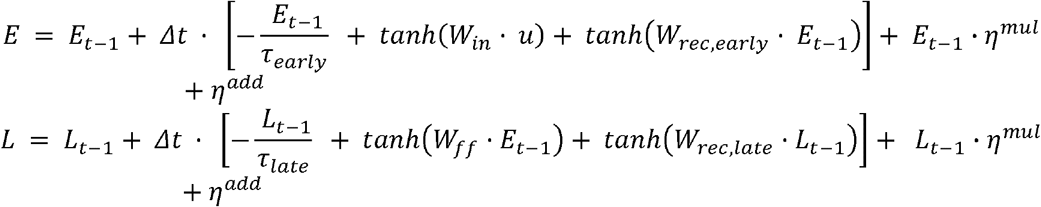

Where:

- *E*_*t*_,*L*_*t*_ are the activity vectors at time *t* for early- and late-regions respectively.
- *W*_*in*_ is the input weight matrix (applied to input stimulus t), delivered only to the early-order layer.
- *W*_*rec,early*_,*W*_*rec,late*_ are recurrent connectivity matrices within each layer.
- *W* _*ff*_ is the feedforward weight matrix from the early to late region.
- *τ* _early_, *τ* _*late*_ are the membrane time constant for each layer.
- *u*_*t*_ is the stimulus vector delivered during the peri-stimulus time window.
- *Δt*is the simulation time step (0.05 s).
- *η*_*t*_ are Gaussian noise terms applied after integration.
  - 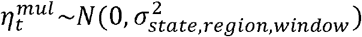 is multiplicative on current activity, with variance determined by the empirical Fano factors computed across the neuronal population for each brain state (wake vs. NREM), layer, and time window (pre-, peri-, post-stimulus).
  - 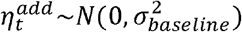is a small additive noise constant (not state- or region-dependent).

Brain state was modeled by adjusting the noise magnitude, with higher variability during wake than sleep, consistent with empirical measurements of the Fano factor and observations of trial-to-trial variability and distractor interference^33^.

The model was tuned to qualitatively replicate observed patterns of stimulus decodability and persistence:

- Early region exhibited similar post-stimulus decay constants across wake and sleep.
- Late region showed prolonged persistence during sleep, in contrast to rapid decay during wake.
- Early region exhibited higher decoding accuracy compared to the late region during stimuli presentation.

Decoding of stimulus identity from the model’s output was performed using the same logistic regression pipeline as the empirical data. Temporal decay constants were extracted via exponential fitting of decoding performance across time as before.

## Acknowledgments

We thank Saar Berkovitch, Anan Moran, and Adi Mizrahi for assistance with Neuropixels setup. Israel Nelken and Ran Darshan for discussions and inputs on manuscript drafts. Supported by grants to Y.N. from the Israel Science Foundation (ISF 1557/22) and the European Research Council (ERC-2019-COG 864353).

## Author Contributions

- Conception and Design of Research: B. Hadad, N. Regev, and Y. Nir.
- Funding Acquisition: Y. Nir.
- Design and Construction of Electrophysiology Experimental Setup: B. Hadad
- Development and Deployment of Chronic Neuropixels device: O. Rechnitz, S. Abramson, and D. Derdikman
- Data Collection: B. Hadad, N. Regev, and U. Kimchi
- Data Analysis: B. Hadad
- Manuscript Writing: B. Hadad and Y. Nir.
- Critical Review of Results and Manuscript Comments: All authors.

## References

1. Nir, Y., Vyazovskiy, V. V., Cirelli, C., Banks, M. I. & Tononi, G. Auditory Responses and Stimulus-Specific Adaptation in Rat Auditory Cortex are Preserved Across NREM and REM Sleep. Cereb. Cortex 25, 1362–1378 (2015).

2. Sela, Y., Vyazovskiy, V. V., Cirelli, C., Tononi, G. & Nir, Y. Responses in Rat Core Auditory Cortex are Preserved during Sleep Spindle Oscillations. Sleep 39, 1069–1082 (2016).

3. Sela, Y., Krom, A. J., Bergman, L., Regev, N. & Nir, Y. Sleep Differentially Affects Early and Late Neuronal Responses to Sounds in Auditory and Perirhinal Cortices. J. Neurosci. 40, 2895–2905 (2020).

4. Marmelshtein, A., Eckerling, A., Hadad, B., Ben-Eliyahu, S. & Nir, Y. Sleep-like changes in neural processing emerge during sleep deprivation in early auditory cortex. Curr. Biol. 33, 2925–2940.e6 (2023).

5. Marmelshtein, A., Lavy, B., Hadad, B. & Nir, Y. Abrupt and gradual changes in neuronal processing upon falling asleep and awakening. J. Neurosci. e1288242025 (2025) doi:10.1523/JNEUROSCI.1288-24.2025.

6. Hayat, H. et al. Reduced neural feedback signaling despite robust neuron and gamma auditory responses during human sleep. Nat. Neurosci. 25, 935–943 (2022).

7. Mariotti, M., Formenti, A. & Mancia, M. Responses of VPL thalamic neurones to peripheral stimulation in wakefulness and sleep. Neurosci. Lett. 102, 70–75 (1989).

8. McCormick, D. A. & Bal, T. Sensory gating mechanisms of the thalamus. Curr. Opin. Neurobiol. 4, 550–556 (1994).

9. Mashour, G. A. Top-down mechanisms of anesthetic-induced unconsciousness. Front. Syst. Neurosci. 8, (2014).

10. Raz, A. et al. Preferential effect of isoflurane on top-down vs. bottom-up pathways in sensory cortex. Front. Syst. Neurosci. 8, (2014).

11. Massimini, M. et al. Breakdown of Cortical Effective Connectivity During Sleep. Science 309, 2228–2232 (2005).

12. Goldman-Rakic, P. S. Cellular basis of working memory. Neuron 14, 477–485 (1995).

13. Curtis, C. E. & D’Esposito, M. Persistent activity in the prefrontal cortex during working memory. Trends Cogn. Sci. 7, 415–423 (2003).

14. Romo, R., Brody, C. D., Hernández, A. & Lemus, L. Neuronal correlates of parametric working memory in the prefrontal cortex. Nature 399, 470–473 (1999).

15. Constantinidis, C. et al. Persistent Spiking Activity Underlies Working Memory. J. Neurosci. 38, 7020–7028 (2018).

16. Lundqvist, M., Herman, P. & Miller, E. K. Working Memory: Delay Activity, Yes! Persistent Activity? Maybe Not. J. Neurosci. 38, 7013–7019 (2018).

17. Curtis, C. E. & Sprague, T. C. Persistent Activity During Working Memory From Front to Back. Front. Neural Circuits 15, 696060 (2021).

18. Harrison, S. A. & Tong, F. Decoding reveals the contents of visual working memory in early visual areas. Nature 458, 632–635 (2009).

19. Romo, R., Hernández, A., Zainos, A., Lemus, L. & Brody, C. D. Neuronal correlates of decision-making in secondary somatosensory cortex. Nat. Neurosci. 5, 1217–1225 (2002).

20. Romo, R. & Salinas, E. Flutter Discrimination: neural codes, perception, memory and decision making. Nat. Rev. Neurosci. 4, 203–218 (2003).

21. Lazar, A., Lewis, C., Fries, P., Singer, W. & Nikolic, D. Visual exposure enhances stimulus encoding and persistence in primary cortex. Proc. Natl. Acad. Sci. 118, e2105276118 (2021).

22. Serences, J. T., Ester, E. F., Vogel, E. K. & Awh, E. Stimulus-Specific Delay Activity in Human Primary Visual Cortex. Psychol. Sci. 20, 207–214 (2009).

23. Huang, Y., Matysiak, A., Heil, P., König, R. & Brosch, M. Persistent neural activity in auditory cortex is related to auditory working memory in humans and nonhuman primates. eLife 5, e15441 (2016).

24. Mesgarani, N., David, S. V., Fritz, J. B. & Shamma, S. A. Influence of Context and Behavior on Stimulus Reconstruction From Neural Activity in Primary Auditory Cortex. J. Neurophysiol. 102, 3329–3339 (2009).

25. Yu, Q., Panichello, M. F., Cai, Y., Postle, B. R. & Buschman, T. J. Delay-period activity in frontal, parietal, and occipital cortex tracks noise and biases in visual working memory. PLOS Biol. 18, e3000854 (2020).

26. Englitz, B., Akram, S., Elhilali, M. & Shamma, S. Decoding contextual influences on auditory perception from primary auditory cortex. eLife 13, RP94296 (2024).

27. Jun, J. J. et al. Fully integrated silicon probes for high-density recording of neural activity. Nature 551, 232–236 (2017).

28. Cavelli, M. L. et al. Sleep/wake changes in perturbational complexity in rats and mice. iScience 26, 106186 (2023).

29. Senzai, Y., Fernandez-Ruiz, A. & Buzsáki, G. Layer-Specific Physiological Features and Interlaminar Interactions in the Primary Visual Cortex of the Mouse. Neuron 101, 500–513.e5 (2019).

30. Vyazovskiy, V. V. & Harris, K. D. Sleep and the single neuron: the role of global slow oscillations in individual cell rest. Nat. Rev. Neurosci. 14, 443–451 (2013).

31. Mazzucato, L., Fontanini, A. & La Camera, G. Stimuli Reduce the Dimensionality of Cortical Activity. Front. Syst. Neurosci. 10, (2016).

32. Churchland, M. M. et al. Stimulus onset quenches neural variability: a widespread cortical phenomenon. Nat. Neurosci. 13, 369–378 (2010).

33. Luczak, A., Bartho, P. & Harris, K. D. Gating of Sensory Input by Spontaneous Cortical Activity. J. Neurosci. 33, 1684–1695 (2013).

34. Musall, S., Kaufman, M. T., Juavinett, A. L., Gluf, S. & Churchland, A. K. Single-trial neural dynamics are dominated by richly varied movements. Nat. Neurosci. 22, 1677–1686 (2019).

35. Esmaeili, V. & Diamond, M. E. Neuronal Correlates of Tactile Working Memory in Prefrontal and Vibrissal Somatosensory Cortex. Cell Rep. 27, 3167–3181.e5 (2019).

36. Atienza, M., Cantero, J. L. & Gómez, C. M. Decay time of the auditory sensory memory trace during wakefulness and REM sleep. Psychophysiology 37, 485–493 (2000).

37. Paluch, K. et al. Unattended working memory items are coded by persistent activity in human medial temporal lobe neurons. Nat. Hum. Behav. (2025) doi:10.1038/s41562-025-02235-0.

38. Strauss, M. et al. Disruption of hierarchical predictive coding during sleep. Proc. Natl. Acad. Sci. 112, (2015).

39. Barak, O., Sussillo, D., Romo, R., Tsodyks, M. & Abbott, L. F. From fixed points to chaos: Three models of delayed discrimination. Prog. Neurobiol. 103, 214–222 (2013).

40. Mante, V., Sussillo, D., Shenoy, K. V. & Newsome, W. T. Context-dependent computation by recurrent dynamics in prefrontal cortex. Nature 503, 78–84 (2013).

41. Nir, Y. & De Lecea, L. Sleep and vigilance states: Embracing spatiotemporal dynamics. Neuron 111, 1998–2011 (2023).

42. Oudiette, D. & Paller, K. A. Upgrading the sleeping brain with targeted memory reactivation. Trends Cogn. Sci. 17, 142–149 (2013).

43. Pachitariu, M., Sridhar, S., Pennington, J. & Stringer, C. Spike sorting with Kilosort4. Nat. Methods 21, 914–921 (2024).

44. Fabre, J. M. J., Van Beest, E. H., Peters, A. J., Carandini, M. & Harris, K. D. Bombcell: automated curation and cell classification of spike-sorted electrophysiology data. Zenodo 10.5281/ZENODO.8172821 (2023).

45. Hayat, H. et al. Locus coeruleus norepinephrine activity mediates sensory-evoked awakenings from sleep. Sci. Adv. 6, eaaz4232 (2020).

46. Rossant, C. et al. Spike sorting for large, dense electrode arrays. Nat. Neurosci. 19, 634– 641 (2016).

47. Ding PhD, M. & Glanzman PhD, D. The Dynamic Brain. (Oxford University Press, 2011). doi:10.1093/acprof:oso/9780195393798.001.0001.

